# Serum Proteomic Profiling Implicates a Dysregulated Neurohormonal-Inflammatory Axis in Post-Fontan Tachycardia

**DOI:** 10.64898/2026.01.30.702962

**Authors:** Felipe Takaesu, Delaney J. Villarreal, Ashley Zhou, Michael R. Jimenez, Mackenzie E. Turner, J. Logan Spiess, Jennifer C. Kievert, Cameron M. DeShetler, William E. Schwartzman, Andrew R. Yates, John M. Kelly, Christopher K. Breuer, Michael E. Davis

## Abstract

**Background:** Post-operative tachycardia is a common and poorly understood complication following the Fontan procedure. Post-operative factors such as surgical scarring and venous hypertension can contribute to tachycardia risk, but the specific molecular signaling cascades triggering acute tachycardia remain uncharacterized, limiting therapeutic innovation and leaving clinicians with only reactive management strategies. Here, we present a retrospective translational study leveraging serum proteomics and machine learning to identify the molecular drivers of post-operative Fontan tachycardia.

**Methods:** We integrated a clinically relevant ovine animal model of the Fontan circulation with continuous telemetric heart rate monitoring and human patient data. Serum proteomics coupled with machine learning algorithms (LASSO and Boruta) were employed to identify protein panels predictive of post-operative tachycardia. Cross-species validation was performed by comparing proteomic signatures from sheep and pediatric patients undergoing Glenn or Fontan surgery.

**Results:** Ovine Fontan animals demonstrated significant heart rate elevation beginning on post-operative day (POD) 1, peaking at POD 3 (159.4 ± 11.7 bpm vs. pre-operative 105.3 ± 10.5 bpm, p<0.0001), before trending toward baseline by POD 10. This pattern was mirrored in human pediatric patients, though with a more modest magnitude. Surgical controls did not exhibit tachycardia, indicating the response was specific to the Fontan procedure. Proteomic analysis identified distinct separation between pre- and post-operative serum profiles. Principal component analysis revealed that the principal components most correlated with heart rate (PC1: r=0.79, p=6.5×10⁻⁶; PC8: r=0.40, p=0.04) were significantly enriched for inflammatory and neural pathways. We leveraged the Boruta algorithm to identify a seven-protein panel (ACE, ANGT, ITIH4, SELENOP, W5PHP7, PTX3, and F5) with superior predictive power (AUC=0.926). A cross-species comparison between human and sheep demonstrated that three proteins, angiotensinogen (ANGT), angiotensin-converting enzyme (ACE), and pentraxin 3 (PTX3), were similarly dysregulated in both species post-operatively.

**Conclusions:** This study provides the first direct molecular evidence implicating a dysregulated neurohormonal-inflammatory axis as a principal driver of acute post-operative Fontan tachycardia. The identified protein signature offers novel mechanistic insights and establishes a foundation for developing targeted diagnostics and therapeutics to predict and mitigate this significant clinical complication.

## Introduction

The Fontan procedure represents the final surgical palliation for patients with single ventricle diseases, a group of congenital heart defects that were once uniformly fatal in early childhood^1,2^. This staged surgical approach transformed patient outcomes, creating an expanding global population of survivors who now reach adolescence and adulthood. Nevertheless, the Fontan operation, while lifesaving, is palliative^3,4^. The Fontan surgery establishes a fragile and abnormal circulation which results in systemic venous hypertension and chronic ventricular preload deficiency^5–7^. This unique hemodynamic state of elevated central venous pressure and relatively low cardiac output is the primary driver of a progressive, multi-system disease burden that defines the long-term clinical course for these patients. The consequences are ubiquitous and severe, including, but not limited to, congestive hepatopathy, protein-losing enteropathy, plastic bronchitis, and renal dysfunction^5,8–10^.

Acute post-operative tachycardia represents a pervasive and poorly understood complication in the immediate period following the Fontan procedure. The underlying causes are often attributed to a combination of factors, such as surgical scarring, the type of Fontan operation, atrial dysfunction as well as the chronic effects of cyanosis and associated end-organ dysfunction^3,11–13^. However, the specific molecular signaling cascades that trigger a tachycardic event, particularly in the vulnerable immediate post-operative period, remain largely uncharacterized. This knowledge gap limits therapeutic innovation and intervention, leaving clinicians with reactive management strategies that often fail to prevent recurrence or address the underlying causes.

Serum proteomics, the large-scale study of circulating proteins, offers a powerful tool to investigate the molecular basis of complex cardiovascular diseases^14^. By directly measuring the proteins that carry out cellular functions, proteomics can uncover disease mechanisms and identify novel biomarkers that other technologies might miss^15,16^. Analyzing the serum proteome provides a unique window into the body’s systemic response, capturing circulating proteins from multiple organ systems that reflect an individual’s physiological state. While previous proteomic studies in Fontan patients identified broad signatures of inflammation, oxidative stress, and metabolic dysregulation, none pinpointed specific molecular pathways directly linked to the onset of tachycardia^17–21^.

In this study, we sought to identify the molecular drivers of acute post-operative tachycardia in the Fontan circulation by integrating a clinically relevant large animal model and human patient data with advanced proteomic and bioinformatic analyses^22^. We leveraged a Fontan surgical ovine model that recapitulates the acute post-operative course and allows for the correlation of high-throughput serum proteomics with continuous telemetric monitoring of heart rate. Using this translational framework, we applied LASSO and Boruta machine learning algorithms to identify protein panels that are highly predictive of post-operative tachycardia in our ovine model. To establish clinical relevance, we confirmed the trend of increased post-Fontan heart rate and performed a comparative proteomic analysis of serum from human pediatric patients undergoing either the Fontan surgery or the immediately preceding procedure, the Glenn, which connects the superior vena cava (SVC) to the main pulmonary artery (MPA). This cross-species validation confirmed that three key proteins from the predictive panel – angiotensinogen (ANGT), angiotensin-converting enzyme (ACE), and the inflammatory marker pentraxin-3 (PTX3) – were similarly dysregulated in human patients post-operatively. This work offers novel mechanistic insight into post-Fontan tachycardias and identifies a panel of candidate biomarkers that may enable the development of future diagnostics and targeted therapeutics to predict and mitigate this condition.

## Methods

### Data availability

The data that support the findings of this study are available from the corresponding author upon reasonable request. Please see the Major Resources Table in the Supplemental Materials for all materials used in this study. Mass spectrometry data used for this study has been deposited to the ProteomeXchange Consortium via the PRIDE partner repository under the dataset identifier: PXD068079.

### Ethical Statement

All animal studies were performed according to ARRIVE Guidelines and under the approval and guidance of the NCH Animal Welfare and Resource Committee (Approval AR13-00079 and AR24-00179). Sheep were housed in facilities that are USDA licensed and AAALAC accredited. Housing space for sheep was in accordance with the 2011 Guide for the Care and Use of Laboratory Animals (including HVAC parameters, lighting, and airflow). Human heart rate measurements were collected and analyzed under an approved IRB (STUDY00000005 and STUDY00000887). Human blood samples were collected under an approved IRB (STUDY00005500) prior to surgery and before discharge from patients undergoing Glenn or Fontan surgery. Informed consent was received from parents prior to study enrollment. This study was performed as a retrospective analysis of collected data and samples.

### Telemetry Device Implantation

Fontan (*n* = 7) and surgical control (*n* = 4) Dorset sheep (*Ovis aries*) were anesthetized and received implanted continuous monitoring telemetry devices (DSI PhysioTel Digital L21-F1 Large Animal Implantable Telemetry Unit). Sheep subject characteristics are detailed in **Table S1**. The procedure was performed as described previously^23^. Briefly, the device itself was placed in a subcutaneous pocket either in the right lower abdomen adjacent to the inguinal crease, or at the base of the left neck between the scapula and cervical spine, while the bipotential ECG leads were placed subcutaneously over the sternum. Pressure transducers were cannulated either in the femoral artery or carotid artery to obtain vitals information. After the implantation, Fontan animals were followed via continuous telemetry monitoring for 10 days at a data collection rate of 6 collections per minute (once every 10 seconds). Surgical control animals were followed for 5 days post-operation in the same manner. Continuous data collected from the telemetry units appears in real-time in the linked Ponemah software which was used to inform intensive care unit (ICU) medical decision making. Collected data was saved and exported to Microsoft Excel for further analysis.

### Establishing the Fontan Connection

The Fontan cohort of ovine subjects underwent an additional surgical operation in a separate anesthetic event at least a month after their telemetry device implantation. This operation established a Fontan connection by detaching the inferior vena cava from the right atrium and reconnecting it to the main pulmonary artery via an end-to-end anastomosis facilitated by a polytetrafluorethylene conduit, as described previously^24^. Following the operation, sheep were closely monitored 24/7 for approximately 2 weeks in an ICU.

### Post-Operation Blood Sample Collection

For both the Fontan cohort and surgical control cohort, blood samples were collected daily from the date of surgery to the end of each animal’s patient-specific ICU monitoring endpoint. Sheep-specific duration of ICU care is noted in **Table S1**. Collected blood samples were utilized for ICU monitoring and for serum proteomic analysis. Samples were clotted using a serum blood collection tube (BD Vacutainer, Cat # 02-683-94) for 15 min at room temperature. Samples were then spun at 1500 × g for 15 min at 4 °C followed by immediate serum collection from the supernatant. Serum samples were stored at -80 °C for long-term use.

Human blood samples were collected under an approved IRB before discharge from patients undergoing either Glenn (n = 5) or Fontan (n = 5) surgery. Samples were clotted using a serum blood collection tube (BD Vacutainer, Cat # 02-683-94) for 15 min at room temperature. Samples were then spun at 1500 × *g* for 15 min at 4 °C followed by immediate serum collection from the supernatant. Serum samples were stored at –80 °C for long-term use. Human patient characteristics for the proteomic analysis are detailed in **Table S2** & **S3**.

### Acute Post-Fontan Dataset Analysis and Statistics

Data analysis of telemetry measurements included pre-operative measurements through POD 10 for the Fontan animals, and the morning after the telemetry implantation operation (POD 1) through POD 5 for the control cohort. Data were separated by day or night according to the 12-hour light/dark cycle of the vivarium with the hours from 6am – 5:59pm being day and 6pm – 5:59am being night. All recorded measurements within this time frame were averaged to generate one value for each POD for each assessed parameter. As there were no statistically significant differences between day and night measurements on any post-operative day (**Figure S1A-C**), and all sample collection was performed during the day, data assessed for the purpose of these study comparisons were the measurements and blood samples collected during the day.

To assess the translatability of our ovine large animal model, we also analyzed human Fontan patient heart rate data for evidence of the same trends (*n* = 26). Fontan patient heart rate data were pulled from patient charts and separated into each POD, as defined as each calendar day after the operation with the date of surgery being POD 0, out to POD 7 as it was the most common date of discharge from the hospital. As this data was obtained from retrospective chart review of the electronic medical record rather than continuous telemetry recording, on occasion, several entries were made within the same hour. In these cases, only the first measurement made for that hour was included in the POD average calculation.

Statistical analysis was performed using GraphPad Prism 10. To assess the significance of each timepoint, we utilized linear regression models with the treatment group as the predictor, or the post-operative timepoint as the predictor, respectively. While Student’s t-tests and Wilcoxon rank sum tests are commonly used, the method of linear regression allows us to obtain the same benefits of these tests for our animal cohorts undergoing repeated longitudinal measurements. We can assess timepoint-specific and overall effect size estimates and obtain the ability to account for the correlated nature of longitudinal measurements and adjust for potential confounders.

### Protein Digestion and Peptide Cleanup

Samples (5 µL) were diluted 1:10 with 8 M urea. Protein reduction was performed using 5 mM dithiothreitol (DTT) for 30 min at room temperature, followed by alkylation with 10 mM iodoacetamide (IAA) in the dark for 30 min at room temperature. Samples were then digested with 4 µg of Lysyl endopeptidase (Wako) overnight. The reaction mixture was then diluted to a final urea concentration of 1 M and subjected to further digestion with 4 µg of trypsin (Thermo Fisher Scientific) overnight at room temperature. Following digestion, the peptide solutions were acidified to a final concentration of 1% formic acid and desalted using 10 mg HLB columns. The columns were conditioned with 500 µL of methanol and equilibrated twice with 500 µL of 1% formic acid. Samples were loaded onto the columns and washed twice with 500 µL of 1% formic acid. Peptides were eluted with 1 mL of 50% acetonitrile and dried to completeness using a centrifugal vacuum concentrator (SpeedVac, Labconco).

### Liquid Chromatography and Mass spectrometry (Vanquish NEO)

Each sample was resuspended in 250 µL of loading buffer containing 0.1% formic acid, and a 1 µL aliquot was analyzed by liquid chromatography coupled to tandem mass spectrometry. Peptide eluents were separated on an Evosep column (8 cm x 150 µm internal diameter (ID) packed with 1.5µm resin) using a Vanquish Neo system (Thermo Fisher Scientific). Buffer A consisted of water containing 0.1% (v/v) formic acid, and buffer B consisted of 80% acetonitrile in water with 0.1% (v/v) formic acid. Elution was carried out over a 16-minute gradient increasing from 1% to 50% solvent B. Peptides were analyzed using an Orbitrap Astral mass spectrometer (Thermo Fisher Scientific) equipped with a high-field asymmetric waveform ion mobility spectrometry (FAIMS Pro) ion mobility source (Thermo Fisher Scientific). A single compensation voltage (CV) of -35 V was applied. Each acquisition cycle began with a full MS1 scan covering a mass range of 380–980 *m/z* at a resolution of 240,000, 500% AGC and 3 ms injection time. Data-independent acquisition (DIA) was performed using higher-energy collision-induced dissociation (HCD) with 2 *m/z* isolation windows covering the full precursor range (380-980 m/z). The total cycle time was set to 0.6 s with a maximum injection time of 2.5 ms. Collision energy was set to 27%, and the scan range was set to 150–2000 *m/z*.

### Database Search

Spectronaut (version 19.9.250512.62635) was used to search all raw files in default library free mode using a human Uniprot database (Downloaded on February, 2019). All parameters were kept at default.

### Proteomic Dataset Pre-processing

Raw protein intensity values from the proteomic analysis were first subjected to quality control. This initial assessment revealed that missing values were predominantly skewed towards proteins with low intensities, suggesting a missing not at random (MNAR) data structure (**Figure S2A**). To address this, missing values were imputed using a K-Nearest Neighbors (KNN) method via the *KNNimp* function from the multiUS R package. Following imputation, the complete dataset was normalized using the normalyzerDE package^25^. Based on the performance evaluation metrics provided by the package, variance stabilizing normalization (VSN) was selected as the optimal method. For subsequent analyses, the human dataset was stratified into separate Fontan and Glenn surgical cohorts.

### Proteomic Data Analysis

Differential enrichment analysis was performed to identify proteins that were significantly altered between pre-operative and post-operative states. For the ovine dataset, the NormalyzerDE R package was utilized to compare the post-operative and pre-operative groups^25^. For the human dataset, differential enrichment results were obtained from a direct comparison between pre-operative and post-operative samples, performed separately for the Fontan and Glenn surgical cohorts. For both species, proteins were considered significantly enriched if they met a threshold of an absolute Log_2_ fold change ≥ 0.48 and an adjusted p-value ≤ 0.05. Adjusted p-values were calculated using the Benjamini-Hochberg method.

Unsupervised analyses were conducted to visualize the overall proteomic profiles. For the human data, these analyses were performed on the combined patient cohort as well as on the stratified Fontan and Glenn groups. Principal component analysis (PCA) was performed using the *prcomp* function in R with scaling and centering applied. Hierarchical clustering was performed on the top 100 most variable proteins using the pheatmap R package. This analysis employed the “ward.D2” clustering method with Manhattan distance and row-wise scaling.

Pathway enrichment analysis was performed on the lists of significantly upregulated proteins from the pre- and post-operative groups using Metascape^26^. For visualization, the top 25 pathways ranked by enrichment score were selected.

### Spearman Correlation and Principal Component Gene Set Enrichment

Principal components of the VSN-normalized ovine proteome were correlated with measured heart rates using Spearman’s rank correlation through the *cor.test* function in R. Biological pathways represented by the significantly correlated principal components were subsequently identified using Principal Component Gene Set Enrichment (PCGSE) analysis through the pcgse package in R^27^. The analysis was run against the Reactome gene set database (c2.cp.reactome.v2024.1.Hs.symbols.gmt), and pathways with a p-value ≤ 0.05 were considered significantly enriched.

### LASSO - and Boruta – based Feature Selection

To identify the most predictive proteins for tachycardia, two feature selection methods were employed: Least Absolute Shrinkage and Selection Operator (LASSO) regression and the Boruta algorithm. The ovine proteomic dataset was first partitioned into training (50%) and testing (50%) sets for model development and evaluation. The LASSO model was implemented using the glmnet R package. The optimal regularization parameter (lambda) was determined using a leave-one-out cross-validation (LOOCV) approach, facilitated by the caret R package. Proteins with non-zero coefficients in the final model were selected as potentially predictive features.

As an orthogonal approach, we used the Boruta algorithm, a random forest-based wrapper. To account for the inherent randomness of the algorithm and ensure a robust feature set, a stability selection protocol was implemented. The Boruta algorithm was run 100 times on the training data. Only proteins that were confirmed as “important” in at least 90% of these runs were retained for the final feature set. For both the LASSO- and Boruta-derived protein lists, the optimal predictive panel was determined by iteratively testing all unique combinations of the selected features. A logistic regression model, using the *glm* function in R, was built for each combination on the training data, and its predictive performance was evaluated on the test data by calculating the Area Under the Receiver Operating Characteristic Curve (AUC) using the *pROC* R package.

## Results

### Ovine post-Fontan tachycardia peaks at POD 3 and begins returning to baseline

Several parameters were assessed from the implanted telemetry devices, including heart rate, systolic blood pressure, and diastolic blood pressure. Only heart rate significantly differed between pre-operation measurements and post-operation, while the other parameters did not change significantly (**Figure S1D and S1E**). Heart rate was found to increase starting immediately after the Fontan operation, with statistically significant differences beginning on POD 1 (p = 0.0007) and continuing to rise on POD 2 (p = 0.0002). The post-operative tachycardia peaked on POD 3 compared to pre-operative heart rates (pre-op 105.32 ± 10.51 bpm, vs post-op 159.40 ± 11.74 bpm, p < 0.0001). Though still maintaining statistical significance, at POD 5, the trend of increasing tachycardia reverses and the difference between pre-operative heart rate and post-operative HR begins to decrease and trend towards a return to baseline (p = 0.0003). This pattern continues as POD 10 measurements continue decreasing in statistical difference compared to pre-operative HR measurements (p = 0.0315) (**Figure 1A**).

**Figure 1.**
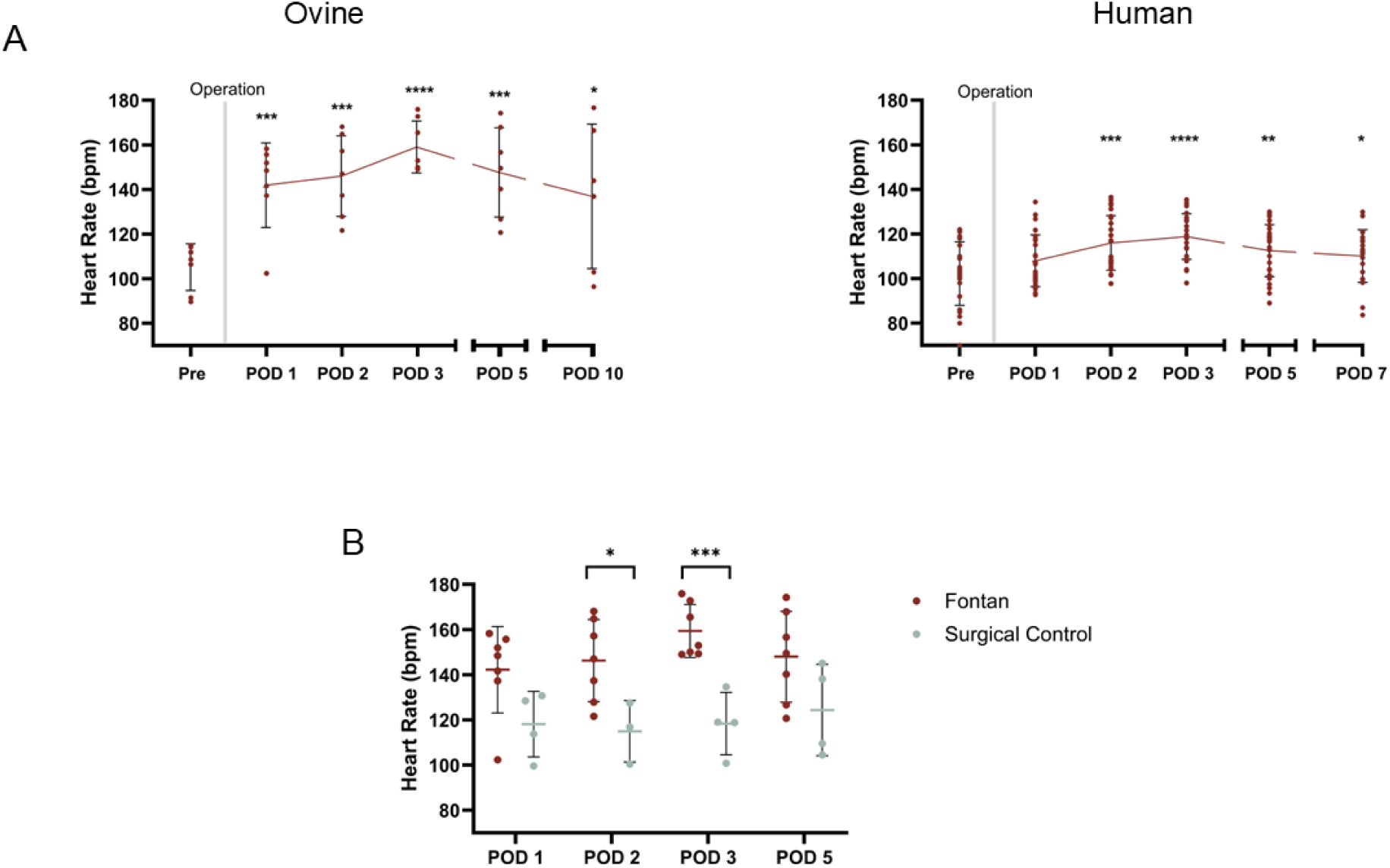
Post-operative heart rate analysis of ovine and human subjects. **(A, left)** Average daily heart rate for each post-operative day (POD) following the Fontan operation in ovine subjects (*n* = 7). Statistically significant increases in average heart rates were observed beginning from POD 1 (\*\*\**p* = 0.0007 | Linear Regression), peaked at POD 3 (*\*\*\*\*p <* 0.0001 | Linear Regression), and began to resolve by POD 5 (\*\*\**p* = 0.0003 | Linear Regression). **(A, right)** Average measured heart rate for each POD following the Fontan operation in human patients (*n* = 26). Patients experienced a statistically significant increase in heart rate beginning at POD 2 following the operation (\*\*\**p* = 0.0005 | Linear Regression), which peaked at POD 3 (\*\*\*\**p* < 0.0001 | Linear Regression), and began to resolve by POD 5 (\*\**p* = 0.0061 | Linear Regression). **(B)** Average daily heart rate following either the Fontan operation (*n* = 7, red line) or a surgical control operation (*n* = 4, blue line) in ovine subjects. Animals receiving the Fontan operation had statistically significant heart rate increases compared to surgical control animals beginning POD 2 (\**p* = 0.029 | Linear Regression), which peaked on POD 3 (\*\*\**p* = 0.0005 | Linear Regression).

Due to the retrospective nature of this data analysis and the intensive care treatment post-operation, animals reaching a heart rate of 160 bpm or greater were administered esmolol. Notably, the ovine subjects were generally symptomatic from the tachycardia. They demonstrated decreased food and water intake as well as reduced physical activity. These clinical signs improved following heart rate control with esmolol. Of the assessed Fontan animals, 6 out of 7 animals required esmolol intervention during their intensive care, with the average number of days on the medication being 3.86 +/- 2.48 days. 4 out of 6 of these animals required esmolol treatment beginning on POD 3. One animal required esmolol beginning on POD 4 and another beginning on POD 6. The longest duration any animal required esmolol intervention during their intensive care treatment was 7 days and the shortest treatment course was 3 days (**Table S1**).

### Pediatric Fontan patients demonstrate a trend of increased post-operative heart rate

The dramatic heart rate increases observed in the ovine Fontan model were compared with trends in human patient data. While no statistically significant differences were observed on POD 1, the pediatric patients did experience a modest heart rate increase beginning POD 2 (p = 0.0005), which peaked on POD 3 (p < 0.0001), before trending towards resolution on POD 5 (p = 0.0061) and POD 7 (p = 0.0492) (**Figure 1A**). The magnitude of observed change was lower compared to the ovine model, with the difference at POD 3 being an average change of 16.7 bpm in humans, compared to 54.08 bpm in sheep. Additionally, no human patients required heart rate modulation with esmolol.

### Ovine surgical controls did not experience post-operative tachycardia

To determine if this post-operative tachycardia was specific to the Fontan operation or generally attributable to surgical insult, the heart rate patterns were compared to a separate cohort of sheep receiving a surgical operation for the implantation of a telemetry monitoring device. This operation consisted of an anesthetic event, incision, device implantation, and wound closure. We compared the heart rates of the Fontan cohort and surgical control cohort on POD 1, 2, 3, and 5, as these corresponded to the rise, peak, and resolution of acute tachycardia in our Fontan operation animals, respectively.

At POD 1, the heart rate of the Fontan animals was not significantly different than the surgical control cohort (p = 0.0579). On POD 3, in contrast to the tachycardia seen in the Fontan cohort, the heart rate of the surgical control cohort remained relatively constant with pre-operative measurements, representing a statistically significant difference between cohorts (p = 0.0005). By POD 5, there was no longer a statistically significant difference in their observed heart rates as the Fontan cohort heart rates begin to return to baseline (p = 0.0942) (**Figure 1B**).

Our analysis supports that our observed post-operative tachycardia is not due to surgical insult alone, as the surgical control cohort animals maintained relatively constant heart rates and significantly lower heart rates than the Fontan animals throughout their recovery. The surgical control cohort’s heart rate measurements did not show the same trend of increasing and resolving tachycardia observed in the Fontan cohort.

As none of the assessed surgical control animals experienced post-operative tachycardia, none required esmolol treatment. This suggests that the post-operative tachycardia observed following the Fontan connection is not driven by anesthetic complications or post-operative pain but rather has a separate etiology.

### The sheep serum proteome is altered by the Fontan operation

To investigate potential molecular mechanisms driving the post-operative tachycardia observed in our sheep, we performed a comparative proteomic analysis of serum collected before and after the Fontan operation. Hierarchical clustering of the top 100 most variable proteins in our dataset revealed a clear separation between pre-operative and post-operative serum profiles (**Figure 2A**). Principal component analysis reinforced that finding with distinct clustering between both sample groups (**Figure 2B**). A scree plot showed principal components one to three representing over fifty percent of the dataset’s variance (**Figure S2B**). Out of the 24 principal components, principal component one (PC1) was the primary driver for clustering as comparisons between other principal components in the absence of PC1 showed significant overlap (**Figure S2C**). Differential enrichment analysis between pre-operative and post-operative groups revealed 104 proteins upregulated in the pre-operative group and 125 proteins being upregulated in the post-operative group (**Figure 2C**).

**Figure 2.**
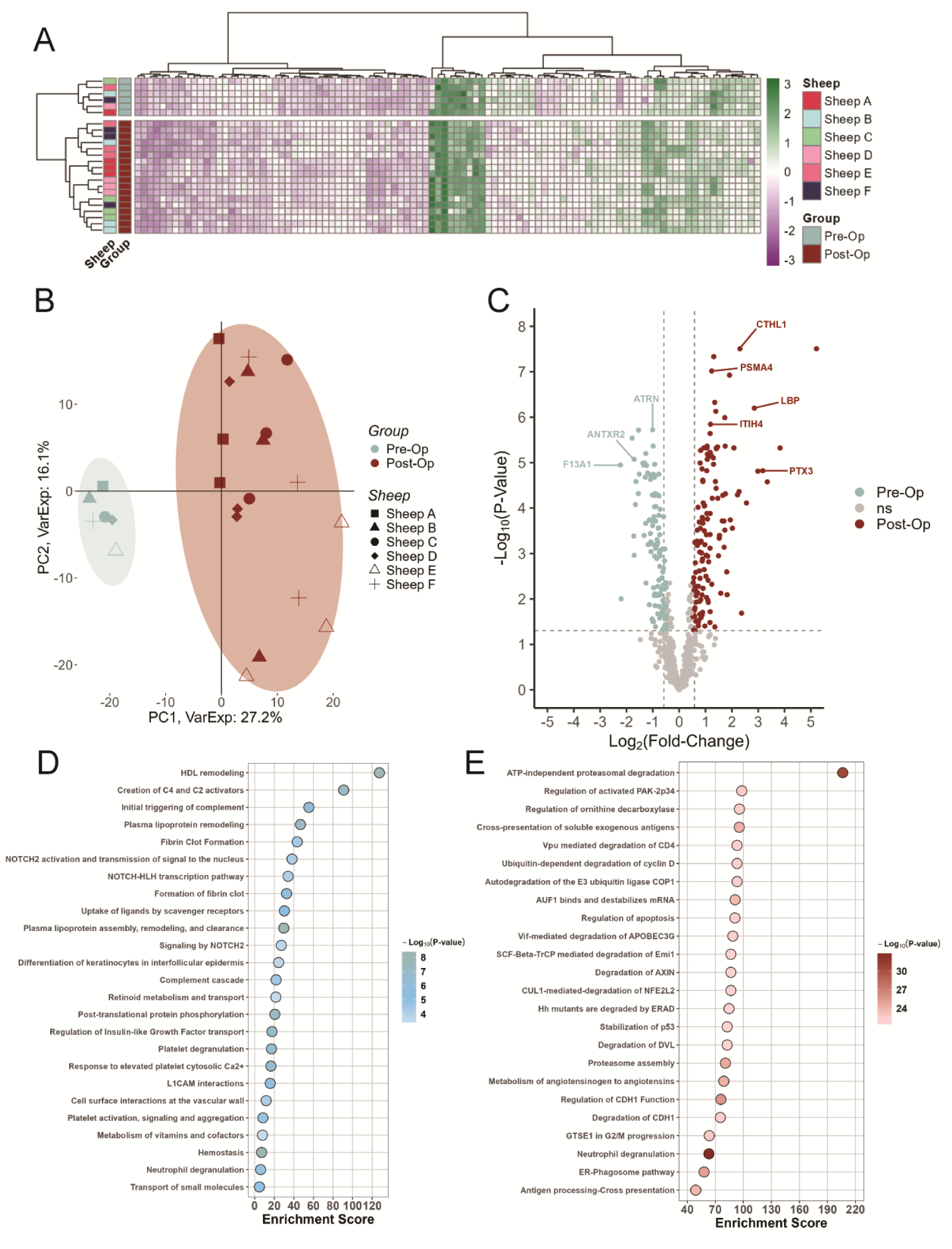
Unsupervised and pathway analysis of the sheep serum proteome pre- and post-Fontan. **(A)** Hierarchical clustering heatmap of the top 100 most variable proteins in our dataset. Clustering for this analysis was done using the “ward.D2” method with Manhattan distance calculations. **(B)** Principal component analysis (PCA) graph of PC1 vs PC2 of the serum proteomic dataset. Shaded areas for the groups were manually added to highlight each group’s respective area. Pre-op (teal, n = 6 with no technical replicates) samples showed clear separation from the post-op (red, n = 6 at different time points) group. **(C)** Volcano plot of the top differentially enriched proteins in a pre-op vs post-op group comparison. 103 and 125 proteins were found to be upregulated in the pre-operative and post-operative groups, respectively. **(D)** Reactome Gene Sets pathway analysis of upregulated proteins found in the pre-operative samples. **(E)** Reactome Gene Sets pathway analysis of upregulated proteins found in post-operative samples.

We next sought to understand the functional implications of these changes by performing pathway analysis to identify biological processes enriched in each group. Pre-operative samples showed an upregulation for pathways predominantly involved in HDL remodeling, the complement system, and hemostasis (**Figure 2D**). The post-operative group was significantly enriched with proteins related to robust inflammatory and protein turnover processes, including neutrophil degranulation and ATP-independent proteasomal degradation, but more interestingly, our results also identified a significant enrichment of the metabolism of angiotensinogen to angiotensin pathway in the post-operative state (**Figure 2E**).

### Inflammatory and neural pathways correlate with post-operative Fontan tachycardia

We next investigated which pathways were most directly associated with the post-operative tachycardia observed in the animals. To achieve this, we correlated the principal components of the serum proteome with heart rates measured at both pre- and post-operative time points. Out of all the principal components, only PC1 (Spearman Cor = 0.79, p-value = 6.5 x 10^-6^) and PC8 (Spearman Cor = 0.40, p-value = 0.04) showed a statistically significant correlation with the heart rate measurements (**Figure 3A** and **3C**). All other principal components had low correlation scores and failed to meet the statistical significance threshold (**Figure S3**). To define the biological functions represented by these principal components, we performed principal component gene set enrichment analysis. This analysis revealed PC1 to be significantly enriched with pathways indicative of a strong inflammatory response, such as innate and adaptive immunity and cytokine signaling, alongside pathways involved in cellular responses to stimuli and nervous system development (**Figure 3B**). PC8 showed a similar enrichment for pathways related to inflammatory responses (**Figure 3D**). A complete list of the PC1 and PC8 loadings can be found on **Table S4** and **S5**. Collectively, these findings suggest that the post-operative tachycardia observed in our sheep Fontan model is driven by a systemic inflammatory state and through neural pathways.

**Figure 3.**
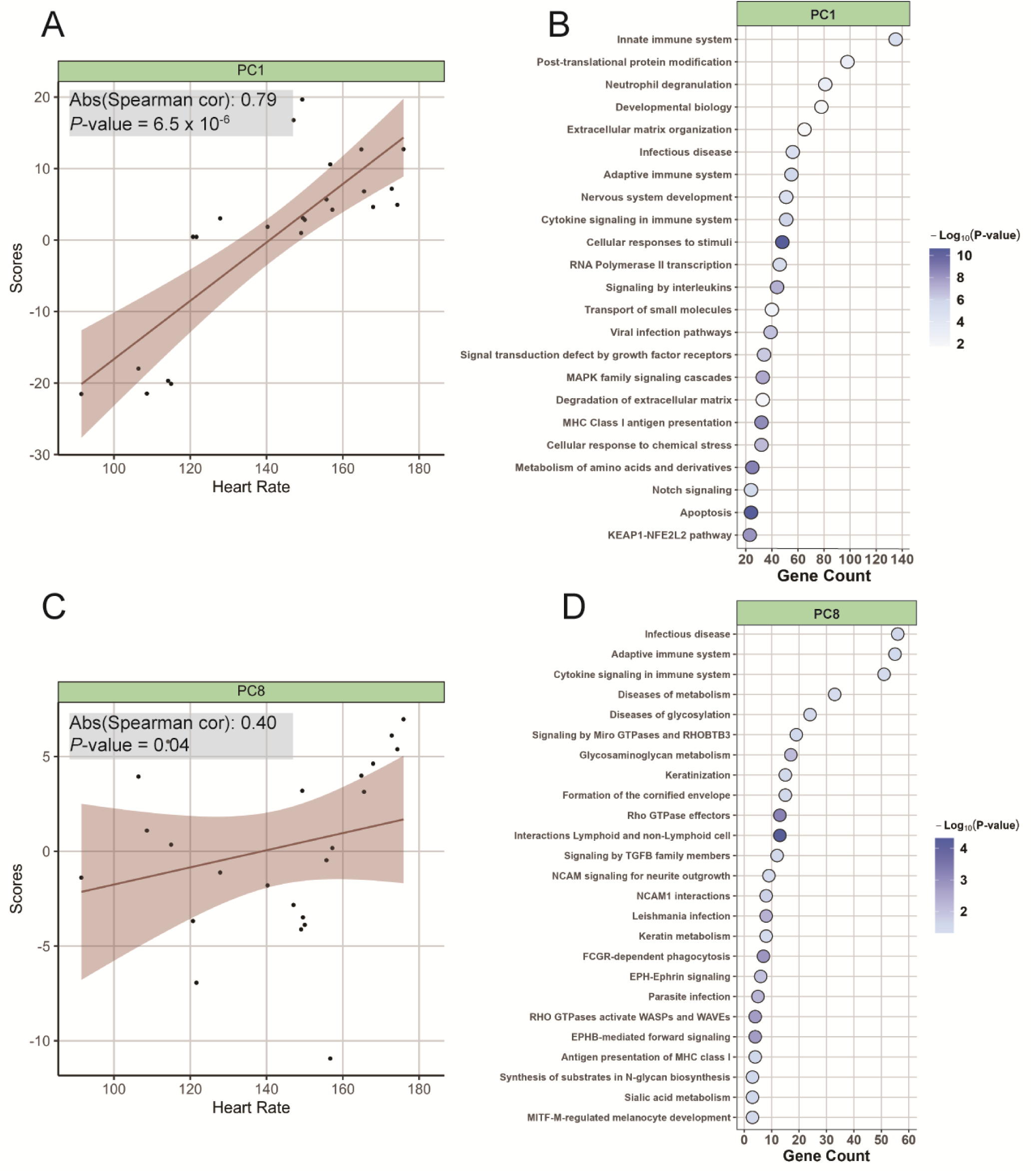
Principal component gene set enrichment analysis of PC1 and PC8 of the sheep serum proteome. **(A)** Spearman correlation (cor) between the first principal component and sheep heart rate measurements. The red shaded area around the best fit line represents the 95% confidence interval. **(B)** Principal component gene set enrichment analysis of the PC1 protein loadings. Pathways were ranked based on their overall gene count in the principal component loadings. **(C)** Spearman correlation (cor) between the eighth principal component and sheep heart rate measurements. The red shaded area around the best fit line represents the 95% confidence interval for the analysis. **(D)** Principal component gene set enrichment analysis of the PC1 protein loadings.

### Identification of key protein drivers of post-operative tachycardia

We then sought to pinpoint the specific proteins found in our pathway analysis that may be mediating the tachycardia. To narrow down the most influential proteins from our dataset, we employed two distinct feature selection methods: Least Absolute Shrinkage and Selection Operator (LASSO) and Boruta (**Figure 4A**). For the LASSO model, we found the optimal regularization parameter to be 0.58 through a leave-one-out cross-validation method (**Figure S4A** and **S4B**). Our final regression with this parameter yielded a final set of 22 proteins as the most predictive features associated with heart rate. To identify the optimal combination of proteins that could be acting as tachycardia drivers, we iteratively tested all 22 unique protein combinations. Our evaluation identified top-performing panels of varying sizes, notably a unique 20-protein combination (AUC = 0.481), a 16-protein combination (AUC 0.500), and an 11-protein combination (AUC = 0.704) that showed the highest predictive potential (**Figure 4B)**. As an orthogonal approach, we utilized the Boruta algorithm to determine if an alternative feature selection method could identify a protein signature with greater predictive power. Surprisingly, the Boruta model identified 9 proteins with strong discriminatory power. By analyzing each protein’s intensity, we found that all nine proteins showed a statistically significant difference in quantity between the pre-operative and post-operative groups (**Figure 4C**). Out of the nine proteins, ITIH4, SELENOP, LBP, and W5PHP7 showed the highest mean variable of importance (**Figure 4D**). After employing a similar iterative combination approach, the seven-protein combination of ACE, ANGT, ITIH4, SELENOP, W5PHP7, PTX3, and F5 showed the highest predictive power (AUC = 0.926) (**Figure 4E**). Finally, the predictive accuracy peaked with this seven-protein panel, as no other combination size achieved a comparable AUC value. These results demonstrate that the Boruta feature selection method identified a superior predictive signature, pinpointing a seven-protein panel (ACE, ANGT, ITIH4, SELENOP, W5PHP7, PTX3, and F5) as potential key drivers of post-Fontan tachycardia in our animal model.

**Figure 4.**
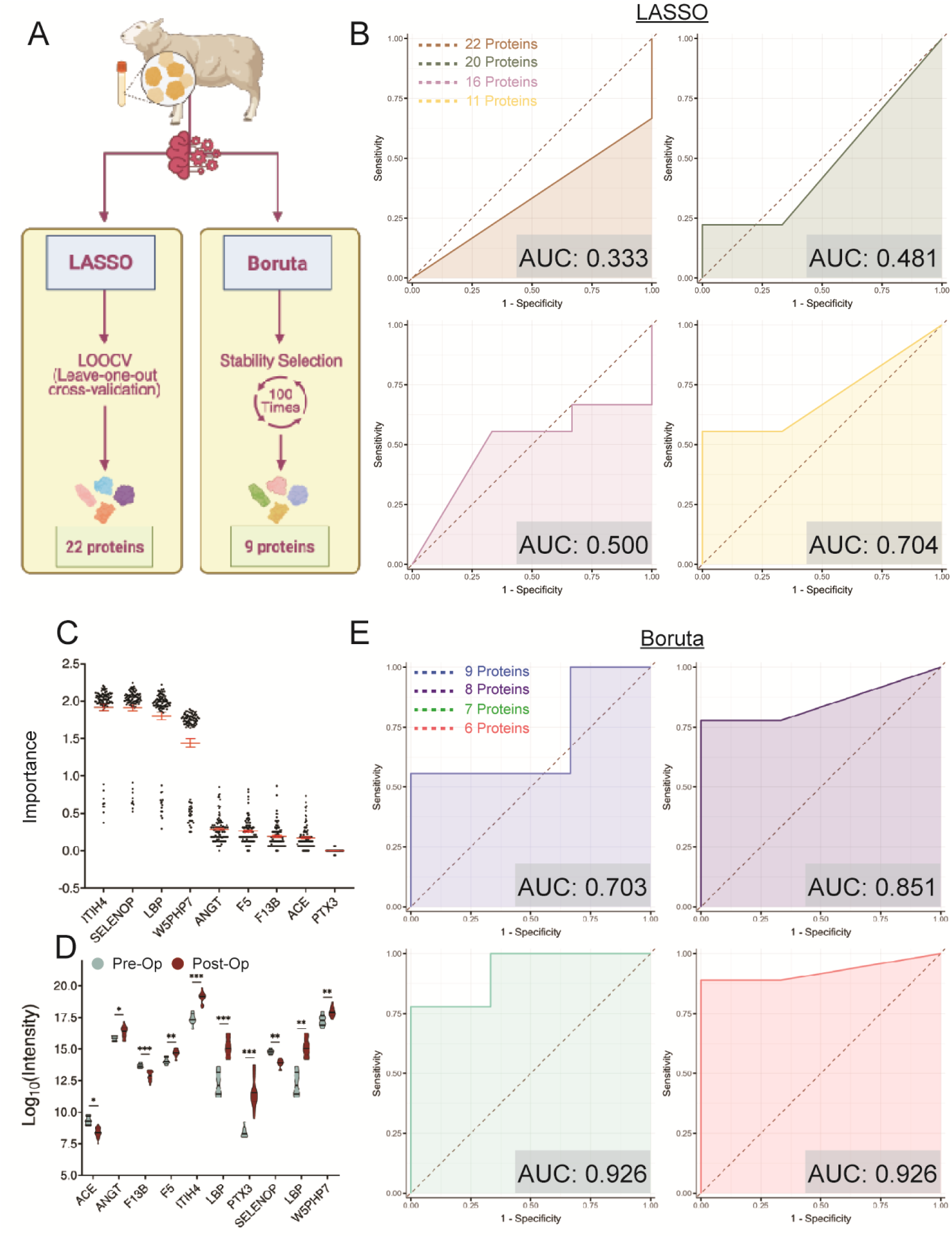
Identification of a candidate protein signature driving post-Fontan tachycardia. **(A)** Graphical representation of our feature selection workflow. To identify the most robust protein signature, two orthogonal methods were employed. First, the Least Absolute Shrinkage and Selection Operator (LASSO) method with leave-one-out cross-validation was used. Second, the Boruta algorithm was applied using a stability selection approach, where the algorithm was run 100 times to identify features consistently ranked as important. **(B)** Receiver operating characteristic (ROC) curves for LASSO protein candidates. The ROC curve plots the true positive rate (sensitivity) against the false positive rate (1-specificity). The diagonal line represents a random classifier with no predictive power (AUC = 0.5). To identify the optimal predictive signature, all unique combinations of the 22 candidate proteins were tested. The curves shown highlight the top-performing combinations, with the shaded area representing the Area Under the Curve (AUC). **(C)** Distribution of mean importance scores for the nine proteins identified by Boruta. The plot illustrates the stability of feature importance across 100 iterative runs of the algorithm. Each point represents the importance value calculated for a given protein from a single run. **(D)** Violin plot illustrating the intensity distribution of the nine Boruta candidate proteins. The plot compares the distribution of protein intensity values between the pre-operative (teal) and post-operative (red) groups for each of the nine candidates. **(E)** Receiver operating characteristic (ROC) curves for the Boruta candidate proteins. The curves plot the true positive rate vs. the false positive rate, where the diagonal line represents random chance. The shaded area indicates the Area Under the Curve (AUC).

### Translational analysis of the human Fontan proteome validates ANGT, ACE, and PTX3 as cross-species markers of post-Fontan tachycardia

To determine the clinical relevance of our findings in the ovine model, we performed a comparative proteomic analysis on serum collected from human patients undergoing either a Glenn or Fontan operation, using a similar pre- and post-operative study design. Hierarchical clustering of the top 100 most variable proteins revealed unclear separation between both groups (**Figure 5A**). A clear separation between pre- and post-operative samples emerged only after stratifying the cohort by surgical stage (Glenn vs. Fontan). This distinct clustering was exclusively observed in the Fontan patient group, while the Glenn patient samples showed no clear separation (**Figure 5B** and **5C**). Further comparisons with different principal components showed similar results with PC1 driving much of the clustering between the pre- and post-operative groups in the Fontan group (**Figure S5**). Differential enrichment analysis also revealed no upregulated proteins in the post-Glenn cohort (**Figure 5D**). On the other hand, 18 upregulated proteins were found in the post-Fontan cohort (**Figure 5E**). We compared the 18 upregulated proteins from the post-Fontan human cohort with the upregulated proteins from our post-operative sheep. This cross-species analysis revealed six conserved proteins: A2GL, ANGT, CATD, G6PI, PGRP1, and PTX3 (**Figure 5F**). Notably, ANGT and PTX3 were also key features identified by our machine learning model as highly predictive of tachycardia. Although ACE narrowly missed the significance threshold for upregulation in the human cohort, its critical association with ANGT and its importance in our ovine predictive model prompted us to investigate it further. Subsequent analysis showed that ANGT, PTX3, and ACE all demonstrated a similar pattern of increased post-operative expression in both human and sheep Fontan groups (**Figure 5G**). Together, these findings suggest that the neurohormonal and inflammatory proteins ANGT, PTX3, and ACE are strong candidates for key drivers of post-Fontan tachycardia.

**Figure 5.**
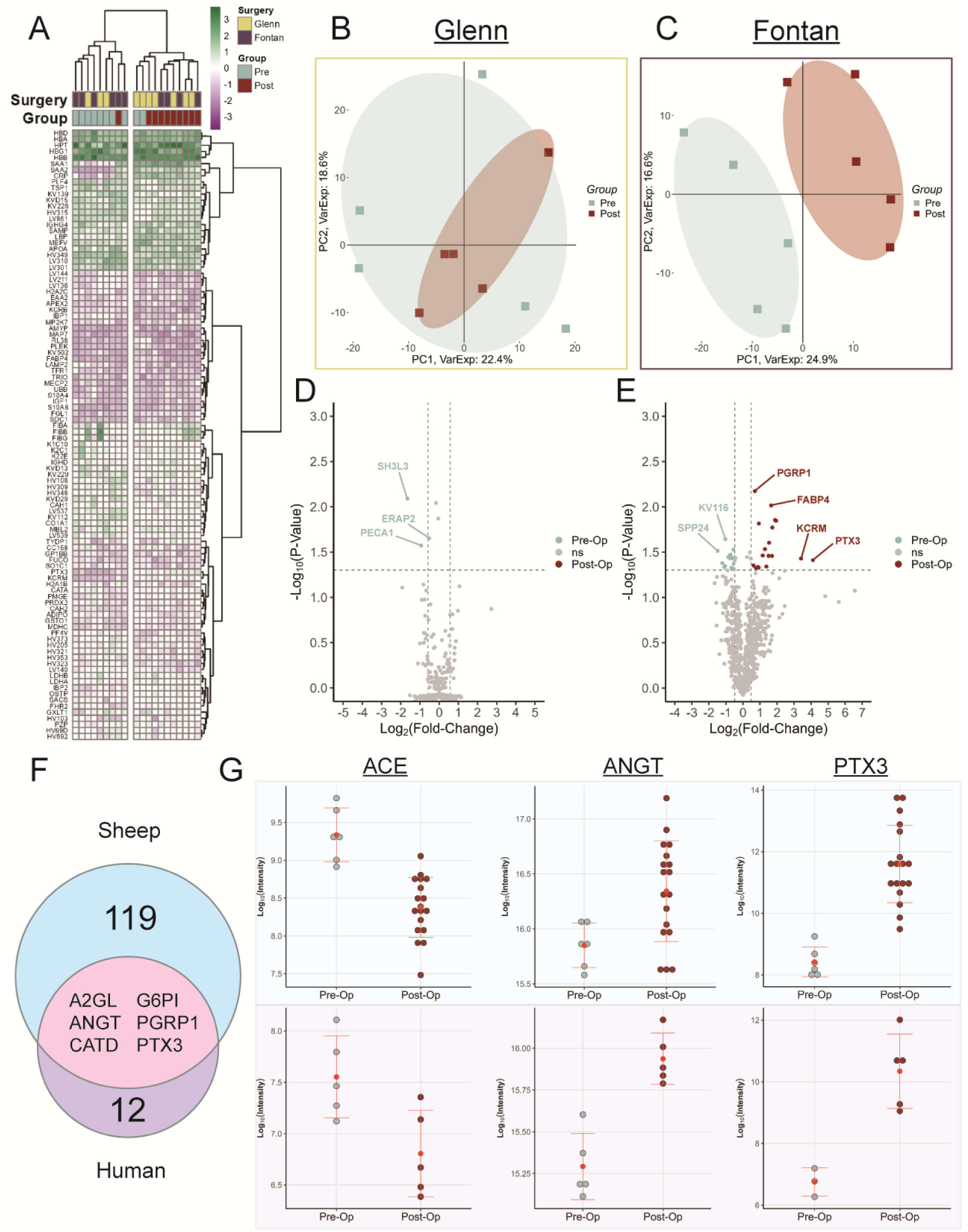
Proteomic profiling of human serum samples pre– and post-Fontan operation. (**A**) Hierarchical clustering of the top 100 most variable proteins in neonatal human serum samples. The heatmap includes samples from both Glenn (yellow) and Fontan (dark purple) patients, annotated by pre-operative (teal) and post-operative (red) time points. **(B)** Principal component analysis (PCA) of the Glenn patient cohort. The plot of PC1 versus PC2 shows no clear separation between pre-operative (teal) and post-operative (red) samples. The shaded area was added manually to highlight the overlapping sample distribution. **(C)** PCA of the Fontan patient cohort. The plot of PC1 versus PC2 shows a distinct separation between pre-operative (teal) and post-operative (red) samples. The shaded areas were added manually to emphasize the clear sample clustering. **(D)** Volcano plot of differential protein expression in the Glenn cohort (n=5). The plot compares post-operative versus pre-operative samples. Proteins with an absolute log2 fold change ≥ 0.48 and an adjusted p-value ≤ 0.05 (indicated by dashed lines) were considered statistically significant. **(E)** Volcano plot of differential protein expression in the Fontan cohort (n=5). The plot compares post-operative versus pre-operative samples. Proteins with an absolute log2 fold change ≥ 0.48 and an adjusted p-value ≤ 0.05 (indicated by dashed lines) were considered statistically significant. **(F)** Venn diagram of upregulated post-Fontan proteins. The diagram illustrates the comparison between upregulated proteins in the ovine (blue) and human (purple) cohorts, revealing a six-protein overlap (pink) between the two groups. **(G)** Cross-species comparison of ANGT, ACE, and PTX3 expression. The plots show the individual protein intensity values for the three key candidates, comparing pre-operative and post-operative samples in both the sheep (blue background) and human (purple background) cohorts.

## Discussion

To our knowledge, this present study provides the first direct molecular evidence implicating a dysregulated neurohormonal-inflammatory axis, specifically centered on the renin-angiotensin-aldosterone system (RAAS), as a principal driver of acute post-operative tachycardia in the Fontan circulation. We first began our investigation through a robust characterization of an ovine Fontan model, which demonstrated high fidelity in replicating the acute post-operative clinical course. The temporal pattern of tachycardia observed in the sheep, with a significant increase beginning on post-operative day one and peaking on day three before trending toward resolution, closely mirrors the timeline observed in pediatric patients recovering from Fontan surgery. This physiological concordance provides confidence that the molecular events captured in the sheep during this period are relevant to humans. The inclusion of a surgical control group further strengthened our findings that the observed tachycardia is not a general response to anesthesia or post-operative pain, but rather a specific pathophysiological response to the unique and abrupt hemodynamic alterations imposed by the Fontan circulation.

The systemic changes induced by the Fontan connection were mirrored by a profound shift in the serum proteome. Unsupervised analyses, including both principal component analysis and hierarchical clustering, revealed an unambiguous separation between pre-operative and post-operative serum profiles in the ovine model. Interestingly, this observation mirrors our previous findings regarding the post-Fontan serum transcriptome^28^. This suggests that the transition to a Fontan circulation, specifically, imposes a systemic insult significant enough to dramatically alter the pre- and post-operative states. An extracellular vesicle deconvolution analysis from our prior work implicated the liver, brain, and other tissue types as primary contributors to the systemic response to the Fontan procedure^28^. Therefore, further experimental work is necessary to deconvolute the complex serum proteome and definitively identify the key tissues driving the proteomic alterations we observe following the Fontan connection.

Having established that the Fontan procedure induces a significant systemic shift in the serum proteome, we sought to elucidate the functional consequences of these alterations. Pathway enrichment analysis of the post-operative proteins revealed a robust inflammatory signature, including the significant enrichment of pathways such as neutrophil degranulation and ATP-independent proteasomal degradation. Intriguingly, this analysis also identified the metabolism of angiotensinogen to angiotensin as a key upregulated pathway, providing the first objective evidence implicating the RAAS system in our animal model. To directly connect these proteomic shifts to the observed tachycardia, we employed a principal component gene set enrichment (PCGSE) analysis. This method identifies the biological processes that are over-represented within the principal components, thereby linking the principal component loadings directly to their biological function. This tool provided a powerful second line of evidence, demonstrating that the principal components most significantly correlated with heart rate (PC1 and PC8) were enriched for pathways related to innate and adaptive immunity, cytokine signaling, and nervous system development. The subsequent identification of ANGT, AGT, and PTX3 as key predictive proteins by machine learning serves as a specific, protein-level validation of this broader pathway-level discovery. The inclusion of PTX3 in this predictive signature is particularly telling. PTX3 can be produced rapidly by cardiac and endothelial cells in direct response to surgical trauma and inflammation, and its elevation is a hallmark of myocardial stress and local inflammation which is known to create a pro-arrhythmic environment by promoting electrical instability^29–33^. Its strong predictive power in our models, therefore, likely represents the initial inflammatory trigger for cardiac irritability^33^.

The specific proteomic signature of the RAAS components in our analysis presents another interesting point of discussion. Previous reports highlight the activation of the RAAS pathway following a Fontan procedure^34^. However, our results show a unique regulatory pattern that warrants further scrutiny. While ANGT, the precursor substrate, was significantly upregulated, we observed a downregulation of ACE in the post-operative group. This pattern deviates from a classic RAAS cascade and points toward an alternative, ACE-independent pathway for angiotensin II (ANG II) generation, likely mediated by the serine protease chymase. In states of profound cardiac stress, previous reports show the heart shifts its primary mechanism of ANG II production to a local, tissue-based chymase-dependent pathway^35,36^. Furthermore, cardiac chymase is not only more efficient at generating ANG II than ACE, but its activity is not inhibited by ACE inhibitors, potentially explaining the persistence of increased heart rate in some patients despite standard therapy^37,38^. Based on our analysis, we suggest a model where the acute surgical inflammation, marked by PTX3, acts as the initial trigger for cardiac irritability, which is then amplified and sustained by the non-canonical RAAS pathway. While an interplay between PTX3 and the RAAS has been reported in other contexts, directly linking this specific inflammatory-neurohormonal axis to the onset of post-operative Fontan tachycardia is a novel insight from our study.

The maintenance of a normotensive state amidst robust RAAS activation represents a significant departure from standard hemodynamics. Because the Fontan circuit lacks a sub-pulmonary pump, it is characterized by chronic preload deficiency and a fundamentally low cardiac output state^39,40^. Under these physiological constraints, the preservation of mean arterial pressure becomes more dependent on compensatory increases in systemic vascular resistance^41^. The upregulation of ANGT identified in our proteomic screen likely provides the substrate necessary to fuel this homeostasis^42^. In this hemodynamic context, normotension may serve as evidence of an active neurohormonal response needed to prevent circulatory collapse.

Mechanistically, we hypothesize that the dissociation between systemic pressure and heart rate can be driven by the enzymatic compartmentalization of the chymase pathway in conjunction with inflammatory modulation by PTX3 (**Figure 6**). Although our data demonstrates systemic ACE downregulation, the high levels of ANGT suggest a shift toward local Angiotensin II production within the myocardium. Previous studies have shown that chymase plays a major role in Angiotensin II formation in the human heart^36,43^. Further literature suggests that chymase activity is strictly localized to the interstitium due to its rapid neutralization by plasma serpins^44^. This biological containment may allow for a local neurohormonal surge that drives tachycardia while the systemic vasculature remains shielded from a hypertensive crisis. Therefore, future studies should look at quantifying the levels of chymase in the cardiac interstitium to investigate this model The identification of PTX3 in our panel adds a layer of inflammatory modulation to this model. Previous studies indicate that PTX3 impairs nitric oxide bioavailability and induces endothelial dysfunction^45^. This mechanism may reinforce the high-resistance state required to sustain blood pressure in the post-operative Fontan heart. Although direct measurement of local Angiotensin II levels was not performed, these proposed mechanisms offer a potential explanation for the divergence between our proteomic signatures and the observed clinical phenotype.

**Figure 6.**
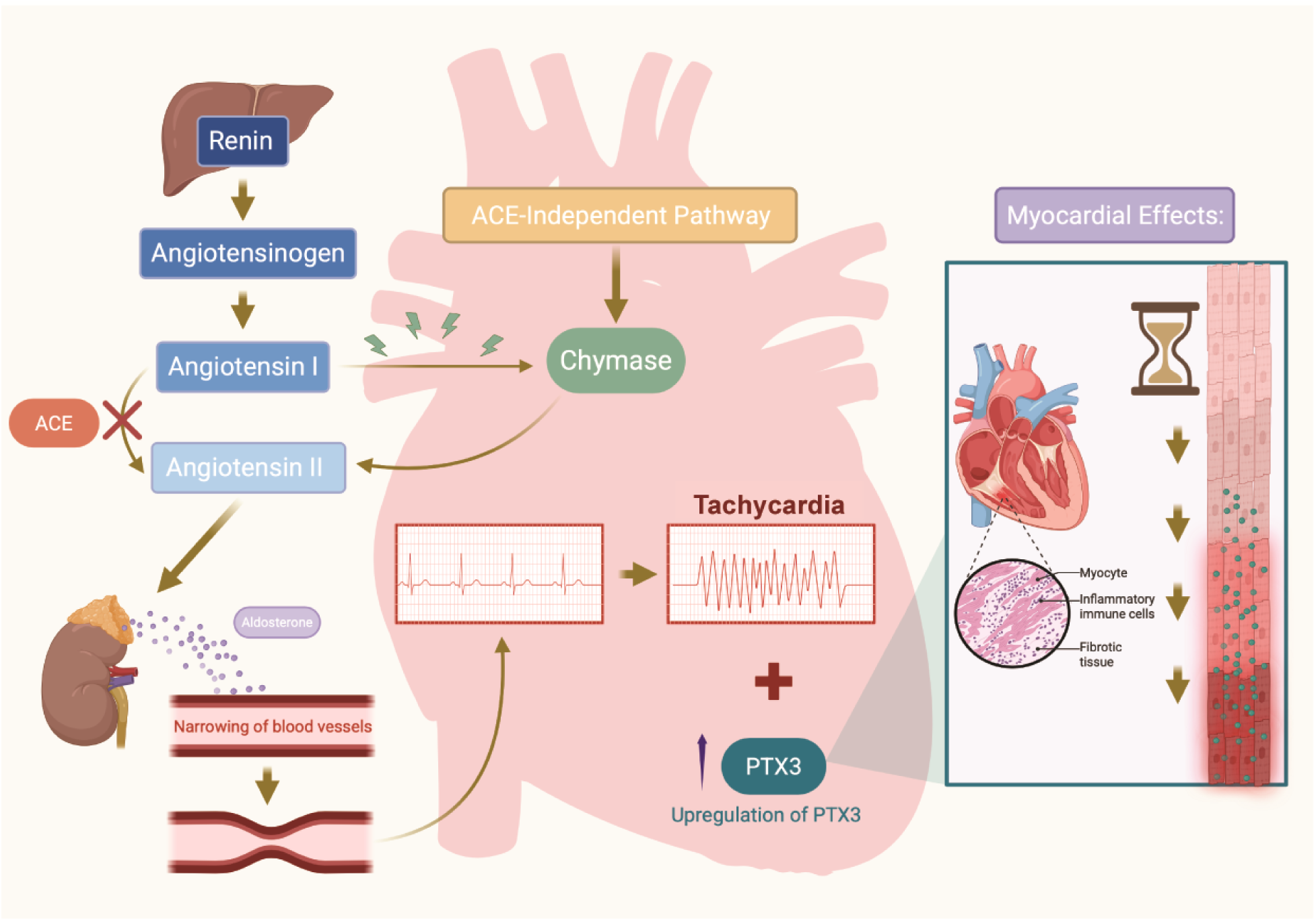
Proposed mechanistic model of the neurohormonal-inflammatory axis driving acute post-operative Fontan tachycardia. The abrupt transition to Fontan hemodynamics triggers a systemic inflammatory response characterized by the upregulation of Pentraxin-3 (PTX3) and the recruitment of inflammatory cells. While systemic Angiotensin-Converting Enzyme (ACE) expression is downregulated, the increase in circulating Angiotensinogen (ANGT) provides a continuous substrate for Angiotensin II production. This process shifts from a systemic ACE-dependent cascade to a localized, myocardial chymase-dependent pathway. The resulting compartmentalized surge in Angiotensin II within the myocardial interstitium promotes cardiac irritability and tachycardia while maintaining necessary systemic vascular resistance through vasoconstriction and aldosterone release.

Finally, this study has some limitations that should be acknowledged. The first is that in humans, Fontan palliation is typically performed in a staged manner. That is, the Fontan operation is commonly performed approximately 2 years following the Glenn operation. However, in the ovine surgical model, Fontan physiology is established in a single stage (i.e. performing the Glenn and IVC to PA extracardiac conduit at the same surgery). As most of the blood flow returning to the heart is derived from the IVC, this abrupt change may be responsible for the more dramatic changes observed in the ovine heart rate data. This is supported further by our proteomic data showing that in human patients, significant differences were only observed after establishing the Fontan circulation and did not appear from the Glenn circulation. Additionally, slight differences exist within the post-operative management that may influence our heart rate and proteomics results. Both our ovine subjects and human patients received spironolactone and furosemide diuretic therapy post-operatively; however, management strategies varied depending on patient needs, duration of treatment and the use of other diuretic medications such as chlorothiazide in humans. Esmolol treatment was not required in human patients because unlike the ovine subjects, which exhibited symptoms from the post-operative tachycardia, the human patients were asymptomatic. Within the animal cohort, the administration of esmolol to treat severe tachycardia was needed in most cases, but not all, and thus introduces a potential confounding variable. The machine learning models were developed on a small cohort, and while the predictive power of the identified protein signature is strong, these findings require validation in a larger animal cohort to confirm their robustness. Finally, the short post-operative follow-up period also limits our ability to assess the long-term molecular signature.

Ultimately, our findings pinpoint ANGT, ACE, and PTX3 signatures as promising, functional readouts of the neurohormonal and inflammatory effectors driving post-Fontan tachycardia. Future therapeutic strategies should target these upstream molecular drivers to mitigate the arrhythmic burden in this vulnerable patient population. Specifically, therapies targeting chymase-dependent Angiotensin II generation or the inflammatory amplification induced by PTX3 may offer novel avenues for clinical management.

## Acknowledgements

We gratefully acknowledge Duc Duong and Kiran Pandey at ARCS Proteomics for their assistance in processing serum samples for mass spectrometry. We also would like to acknowledge the Animal Resources Core at Nationwide Children’s Hospital for the care of the animals used in this study.

## Author Contributions

F. Takaesu, DJ. Villarreal, CK. Breuer, and ME. Davis conceptualized the study. DJ. Villareal, MR. Jimenez, ME. Turner, LJ. Spiess, JC. Kievert, CM. DeShetler, JM. Kelly, and CK. Breuer performed sheep telemetry measurements and analysis. DJ. Villarreal and AR. Yates provided human patient telemetry measurements and analysis. F. Takaesu and A. Zhou performed proteomic and statistical analysis. F. Takaesu, DJ. Villarreal, and A. Zhou created figures and wrote the original draft of the manuscript. All authors agreed with the final version of the manuscript.

## Supplemental Material

Figure S1 – S5

Tables S1 – S5

Major Resources Table

## Novelty and Significance

### What is Known?

- Post-operative tachycardia is a pervasive complication following the Fontan procedure that contributes to significant morbidity in single-ventricle patients.
- Current clinical understanding attributes this complication to non-specific factors such as surgical scarring or atrial dysfunction rather than defined molecular signaling cascades.

### What New Information Does This Article Contribute?

- This study identifies a conserved neurohormonal-inflammatory axis involving angiotensinogen, angiotensin-converting enzyme, and pentraxin-3 that drives acute post-Fontan tachycardia.
- Proteomic profiling suggests the activation of a non-canonical Renin-Angiotensin-Aldosterone System pathway that may be mediated by chymase rather than systemic ACE.
- Machine learning analysis of serum proteomics successfully isolates a specific protein panel that predicts the onset of tachycardia with high accuracy.

Post-operative tachycardia following the Fontan procedure remains a significant clinical challenge with a poorly defined etiology. This study provides the first direct molecular evidence linking a dysregulated neurohormonal-inflammatory axis to this complication. By integrating a high-fidelity ovine model with human clinical data, we identified a specific proteomic signature characterized by the upregulation of angiotensinogen and pentraxin-3 alongside the downregulation of systemic ACE. These findings challenge the traditional understanding of post-operative tachycardia by implicating a non-canonical Renin-Angiotensin-Aldosterone System pathway that is likely driven by local chymase activity. The application of machine learning to serum proteomics also establishes a robust panel of biomarkers with high predictive power. This translational work shifts the paradigm from reactive management to potential targeted therapeutic strategies that address the upstream molecular causes of Fontan-associated tachycardia.

## Non-standard Abbreviations and Acronyms

ANGT: Angiotensinogen
ACE: Angiotensin-converting enzyme
PTX3: Pentraxin 3
POD: Post-operative day
Pre-Op: Pre-operative
Post-Op: Post-operative
PC: Principal component
PCGSE: Principal component gene set enrichment
AUC: Area under the curve
LASSO: Least absolute shrinkage and selection operator
KNN: k-nearest neighbors
VSN: Variance stabilizing normalization
MNAR: Missing not at random
RAAS: Renin-angiotensin-aldosterone system

